# A slow dynamic RNA switch regulates processing of microRNA-21

**DOI:** 10.1101/2021.12.07.471640

**Authors:** Matthew D. Shortridge, Wen Yang, Matthew J. Walker, Gabriele Varani

## Abstract

The microRNAs are non-coding RNAs which post-transcriptionally regulate the expression of a majority of eukaryotic genes, and whose dysregulation is a driver of many human diseases. Here we report the discovery of a very slow (0.1 sec) conformational rearrangement at the Dicer cleavage site of pre-miR-21 which regulates the relative concentration of readily processed and inefficiently processed structural states. We show this dynamic switch is affected by single nucleotide mutations and can be biased by small molecule and peptide ligands, which can direct the microRNA to occupy the inefficiently processed state and reduce processing efficiency. This result reveals a new mechanism of RNA regulation and suggests a chemical approach to suppressing or activating pathogenic microRNAs by selective stabilization of the unprocessed or processed state.

## Introduction

The canonical microRNA (miRNA) biogenesis cascade is a highly orchestrated and regulated process that generates the mature functional single stranded miRNAs from much larger primary transcripts (pri-miRNAs) and intermediate precursors (pre-miRNAs)^1-3^. Recent cellular kinetic studies have demonstrated quantitatively that a precise regulation of miRNA biogenesis is critical for cellular homeostasis^4^; furthermore, dysregulated expression of many microRNAs has been causally associated with human disease since their early association with human cancer^5^, yet it remains unclear how different miRNA precursors are selected for processing. MicroRNA processing enzymes are limiting in the cell and must select amongst a pool of at least 1,500 unique but structurally closely related microRNA precursor species. Changes in expression of specific miRNAs in response to extra- or intracellular stimuli lead in many cases to the development of chronic disease, and suppression of microRNA activity using anti-sense chemistry has shown that disease can be redressed, at least in cellular and animal models^6-8^. However, the limited clinical effectiveness of these agents has prompted efforts to use small molecule chemistry to target the processing enzymes (Drosha/Dicer), ancillary processing proteins (TUTase) or the RNA precursors themselves^9, 10^. Clearly, the discovery of such small molecules, and a basic understanding of microRNA biology, requires better knowledge of the mechanisms by which microRNA expression is regulated, to facilitate identification of targetable steps and structures in the microRNA biogenesis pathway.

Multiple efforts have been dedicated to understanding how the processing enzymes recognize stem-loops common to both primary and precursor microRNAs that encompass the mature miRNA sequences, but not unrelated RNA structures11, 12 (Figure 1A). Enzymatic processing steps are executed by the microprocessor complex with Drosha-DGCR8 in the nucleus and the miRNA Induced Silencing Complex miRISC in the cytoplasm13-20. Dicer is an RNAseIII enzyme which cleaves both dsRNA and pre-miRNA hairpins efficiently and non-specifically (Figure 1B). The miRISC recognizes both the apical loop and the 3’- end of the RNA to orient the enzyme on the substrate and generate the mature 20-22 nt single stranded sequence. It is composed of the RNA binding proteins (RBPs) Dicer, TAR RNA Binding Protein (TRBP), and Argonaute 2 (Ago2). TRBP contains three dsRNA binding domains and improves Dicer processing of pre-miRNAs relative to dsRNA fragments, thereby imparting some specificity. These enzyme complexes are in turn regulated by RNA binding proteins which, in several well-studied examples, bind to the common apical loop structure found in both the pri- and pre-miRNAs21-25.

**Figure 1.**
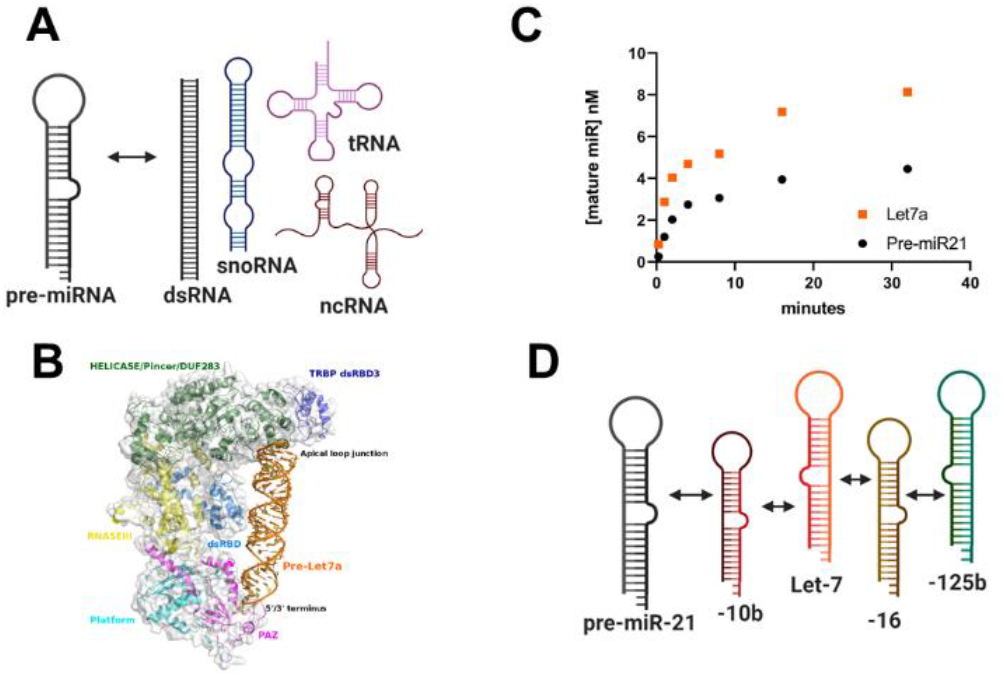
(A) The microRNA processing enzymes distinguish pre-miRNA stem-loop structures from unrelated cellular RNAs. (B) The mechanism for selective cleavage of pre-miRNAs was revealed by recent cryo-EM structures of the Dicer-pre-miRNA complex^20^, but (C) different pre-miRNAs are processed at different rates, as exemplified here by Let-7 and miR-21. Here we show that the slow processing rate of pre-miR-21 is associated with a very slow (approximately 10 Hz) conformational rearrangement of the 5’ dicer cleavage site, which is observed in some microRNAs (e.g. miR-10b) but not others (e.g. pre-Let-7) (D).

Despite being processed by the same enzymes and sharing very similar RNA structures, different pre-miRNA sequences are processed with different efficiency in vitro^11, 26^ and in the cell^4, 15^. For example, the tumor suppressor pre-Let7 is processed nearly five-fold more efficiently than pre-miR-21 by Dicer, and 11-fold more efficiently by Dicer-TRBP^11, 26^ (see also Figure 1C). Mechanistic explanations for these differences have yet to emerge; for example, binding affinities (Kds) for both pre-Let-7 and pre-miR-21 towards Dicer and Dicer-TRBP are very similar^11, 26^.

Many miRNA precursors, including pre-miR-21 (Fig. S1) have a large open apical loop devoid of stable base pairs, that provides a binding site for RNA-binding regulatory proteins and the DGCR8 component of the microprocessor complex^21, 22, 27, 28^. It is likely that local structural and dynamic features of microRNA precursors contribute to making them more or less efficiently processed^29^, but how this occurs remains to be fully understood at the atomic and molecular levels (Figure 1D). Acquisition of this knowledge has been limited by the dynamic properties of both the miRNA substrates and processing proteins^30^; even recent cryo-EM structures have not resolved structural details of the apical loop^17-19^, which is a critical site for regulation by cellular factors.

We and others reported that the apical loop of pre-miR-21 is unstructured and highly dynamic^15, 31, 32^ (Fig. S1), while the Dicer cleavage site is well ordered in free pre-miR-21 but in a dynamically disordered conformation when bound to a peptide macrocycle^31^. We now report that, in the full pre-miR-21 of 59 nts, but not in shorter model fragments studied in the past^33, 34^, the Dicer cleavage site exists in two conformations that exchange with very slow kinetics, at a rate of approximately 10 Hz, corresponding to bulged-out and stacked-in structures of nucleotide A29 at the Dicer cleavage site. By analyzing a set of mutations, we show that the two different structural states represent efficiently (‘on’) or inefficiently (‘off’) processed states and the conformational rearrangement between these two states likely contributes to the slow processing rate of pre-miR-21. We observe multiple conformational states in another slowly processed microRNA, miR-10b, but not in a miRNA which is efficiently processed, Let-7a, suggesting that Dicer prioritizes substrates by selecting those with a pre-existing ‘on’ conformation. Lastly, we show that peptidic and small molecule ligands which bind to pre-miRNA-21 bias the conformational equilibrium towards the non-processed state, thereby reducing processing rates even if they bind to the opposite (major groove) face of the RNA from Dicer, which executes the cleavage chemistry in the minor groove.

## Results

### Very slow conformational dynamics at the Dicer cleavage site of pre-miR-21

Several NMR studies of structure and dynamics of pre-miR-21 utilized short hairpin models to facilitate the spectral analysis^33, 34^. Recently, a conformational switch associated with protonation of A29 at the 5’-Dicer cleavage site and formation of an A+-G base pair was proposed from NMR studies of such a short RNA^34^; we have also long observed the characteristically shifted and slowly exchanging NH2 resonances from A29 associated with formation of the protonated base pair (Figure S2). However, the functional unit recognized by Dicer-TRBP is a 59 nt pre-miR-21 stem-loop (Figure 2A), not these much shorter models, which have no biochemical or biological role. Superficially, the ^1^H-^1^H NOESY spectra of the two miR-21 stem-loop constructs are very similar (Fig. S2); both sequences share the same large and unstructured loop (nucleotides U31-G44)^15, 31, 32^; the putative tandem U31:G44/G32:U43 wobble pairs predicted by computational folding were unobservable in both RNAs as well, only transiently formed at low temperature and unstable (labeled with blue dashes in Figure 2A).

**Figure 2.**
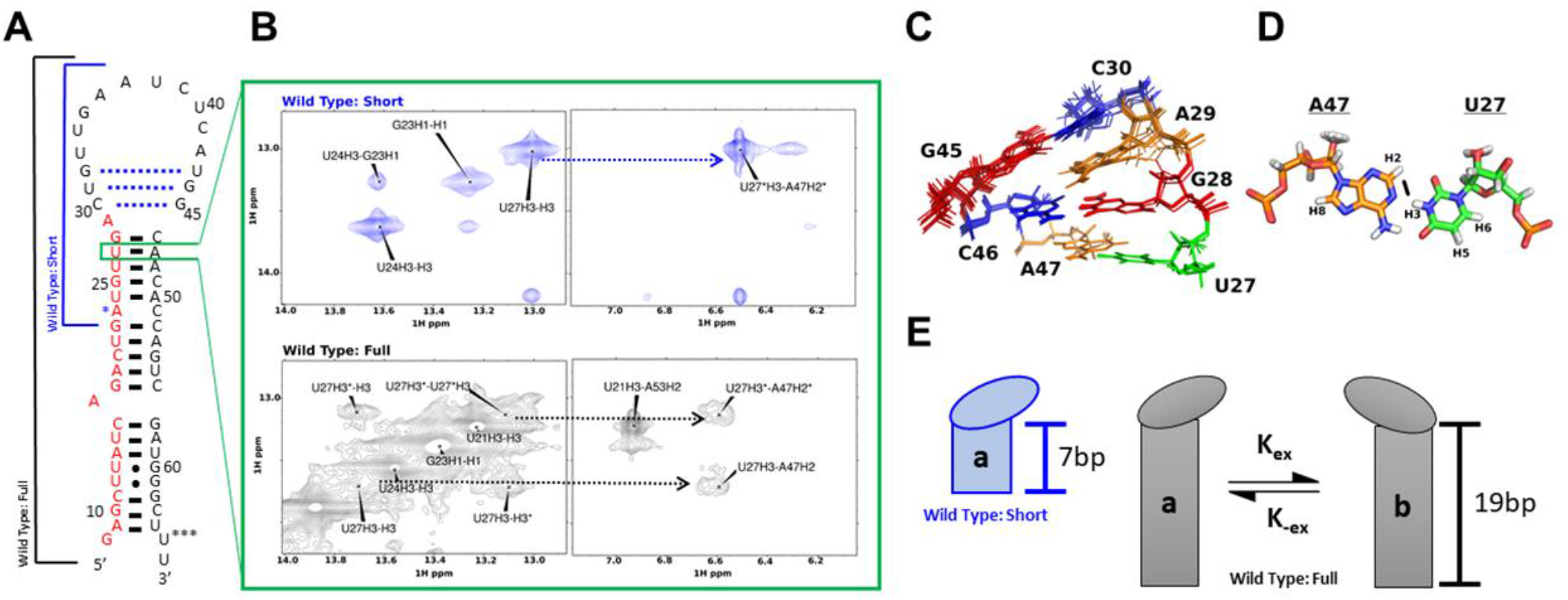
A very slow conformational switch at the Dicer cleavage site of pre-miR-21. A.) NMR derived secondary structure of pre-miR-21; the short miR-21 model used by us and others is boxed in blue (nts G22-C52); the actual (G8-U66) premiRNA generated by Drosha-DGCR8 cleavage and recognized by Dicer-TRBP is in black. For the short sequence, an A23G change was used to improve transcription (*); similarly, the G8-U65 pair (***) was swapped to GC to improve transcription in the full pre-miR-21. B.) The shorter construct does not accurately reproduce the dynamic switch observed in the complete premiR-21. U27 undergoes chemical exchange in the full length pre-miR-21, with two resonances for U27 NH having the same NOE interaction with A47 H2, indicative of base pairing, but this is not observed in the shorter RNA, where a single cross-peak is observed instead. All spectra were collected at 800 MHz and 0.5 mM RNA concentration in 10 mM phosphate buffer pH 6.5, plus 10 mM NaCl, 100 μM EDTA, and at 5C. C.) The pre-miR-21 Dicer cleavage site junction of the short hairpin shows a stacked-in A29 nucleotide, with distinctively shifted NH2 resonances (Figures S1 and S2) consistent with formation of a protonated A+-G base pair^34^; but this nucleotide experiences slow conformational exchange in the full pre-miR-21. D). The distinctive close contact between U27 H3 and A47 H2 from the U27-A47 base pair (black bar) provides a well resolved NOE signal to observe conformational exchange at the pre-miR-21 Dicer cleavage junction. In the short hairpin model, this signal is a singlet (B, top spectra in blue), but in the actual pre-miR-21, it is a doublet (B, bottom spectra in black), each signal corresponds to a distinct conformation of the neighboring residues. E.) Schematic representation of the A29 base flipping mechanism responsible for the two magnetic environments resulting in the chemical exchange peaks observed on U27 and G28.

However, an in-depth analysis of the NMR spectra revealed significant differences at the Dicer-cleavage site (residues G28-A29-C30 and G45-C46). In the 59 nt pre-miRNA that is the actual substrate for Dicer-TRBP and not just a convenient model, the characteristically shifted NH2 resonances from the protonated A29 are not visible at all (Figure S2). In addition, the U27 NH resonance, which is a single peak in the 31 nt short NMR models (Figure 2B, blue), is very clearly split into two peaks separated by 480 Hz (at 800 MHz), revealing the presence of two conformations exchanging at rates much slower than ms (Figure 2B, black; the peak splitting was only observed clearly at high magnetic field strength because of weak intensity and closeness to the diagonal). This slow conformational exchange is also observed in an intermediate stem-loop model of pre-miR-21 (see below), which retains a full helical turn below the Dicer cleavage site (Fig. S3), but not in the shorter hairpin model used in other investigations of pre-miR-21 dynamics^34^.

Exchange rates between the two exchanging conformations were calculated from ^1^H-^1^H EXSY spectra with mixing times of 10 and 200 ms. Peak volumes for the diagonal and off diagonal resonances for the U27 NH were measured directly using SPARKY^35^ and imported into EXSYCALC (Mestrenova) to generate the exchange rate matrix, from which the two rate constants k and k-1 were extracted. The two chemical shifts of U27 originating from distinct magnetic field environments correspond to two comparably populated conformations, U27 and U27*, exchanging with rates: U27→U27* 6.1 s-1 and U27*→U27 7.4 s-1, which corresponds to flip rates of about 10Hz. This rate is approximately 100 times slower than the dynamics observed in the short model hairpin^34^, and suggests the presence of a significant rearrangement in the RNA secondary structure at or near the Dicer cleavage site.

### A29 flipping in and out of the helix reorganizes the structure of the Dicer cleavage site

Despite the presence of two distinct chemical shifts for the U27 NH, this base remains base paired to A47 in both conformations, as demonstrated by the presence of strong cross peaks to the cross strand A47 H2 resonance (at 6.6 ppm) for both conformations (Figure 2B). This observation implies that the U27-A38 base pair does not actively participate in the conformational exchange, but that its resonances are indirectly affected by the rearrangement of nearby residues instead.

To identify the structural mechanism responsible for chemical exchange of U27 NH, we re-examined existing information on the structure of pre-miR-21. The first deposited structure (PDB: 2MNC) concerned a mutated and shortened apical loop, in which the tandem U31:G44/G32:U43 wobble pairs were stabilized to C31:G44 and G32:C43, respectively^33^. For this artificial sequence, the authors noted clear NOEs between G28 NH and G45 NH protons and concluded that A29 was in the ‘bulged-out’ conformation suggested by computational secondary structure predictions. We observed the opposite situation for the short but fully wild-type hairpin model we investigated (PDB 5UTZ)^31^, where A29 adopted a ‘stacked-in’ conformation (Figure 2C/D), wherein the base is at least transiently protonated and base paired^34^, as revealed by well resolved NOEs between the characteristic amino protons of A29 and the imino protons of G28 (11.6 ppm) and G45 (12.5 ppm), along with sequential and cross-strand NOEs between A29 H2, C30 H1’ and C46 H1’.

These observations suggest that A29 can occupy both conformations, bulged outside of the helix or stacked-in; these two conformations could very easily be associated with large chemical shift differences, as observed for U27 NH, because of ring current effects. If the base flipping of A29 between bulged-out and stacked-in conformations was responsible for the chemical exchange reported by the U27 NH chemical shift, we would expect to see split resonances for G28 and probably G45 as well. However, we were unable to identify the corresponding imino proton resonances in the full pre-miR-21 of 59 nts, despite exhaustive efforts. Therefore, we designed an intermediate miR-21 hairpin which includes a full double helical turn below the Dicer cleavage site (nts G18 to C56) and found this sequence to reproduce the dynamic behavior of the full-length construct (Figure 3A). This ‘intermediate’ 39 nt sequence has reduced spectral overlap and improved peak shapes compared to the full pre-miR-21, as expected. Although we still could not observe G45 NH, the improved spectral quality allowed us to observe G28 and measure the exchange rates for both U27 and G28 (Figure 3B), which were U27→U27* 10.7 s-1 and U27*→U27 9.0 s-1 and G28*→G28 (4.6 s-1) and G28→G28* (6.0 s-1). As with the full pre-miR-21 sequence, we observe nearly symmetric exchange between the two states, corresponding to nearly equal populations exchanging at rates of about 10 Hz. The differences in absolute exchange rates between the full length and intermediate RNA models are likely due to differences in overall molecular tumbling rates, as well as uncertainties in experimental measurements, but are obviously very close in magnitude. For this intermediate pre-miR-21 model, the two conformations coalesce at 37C (Figure 3C).

**Figure 3.**
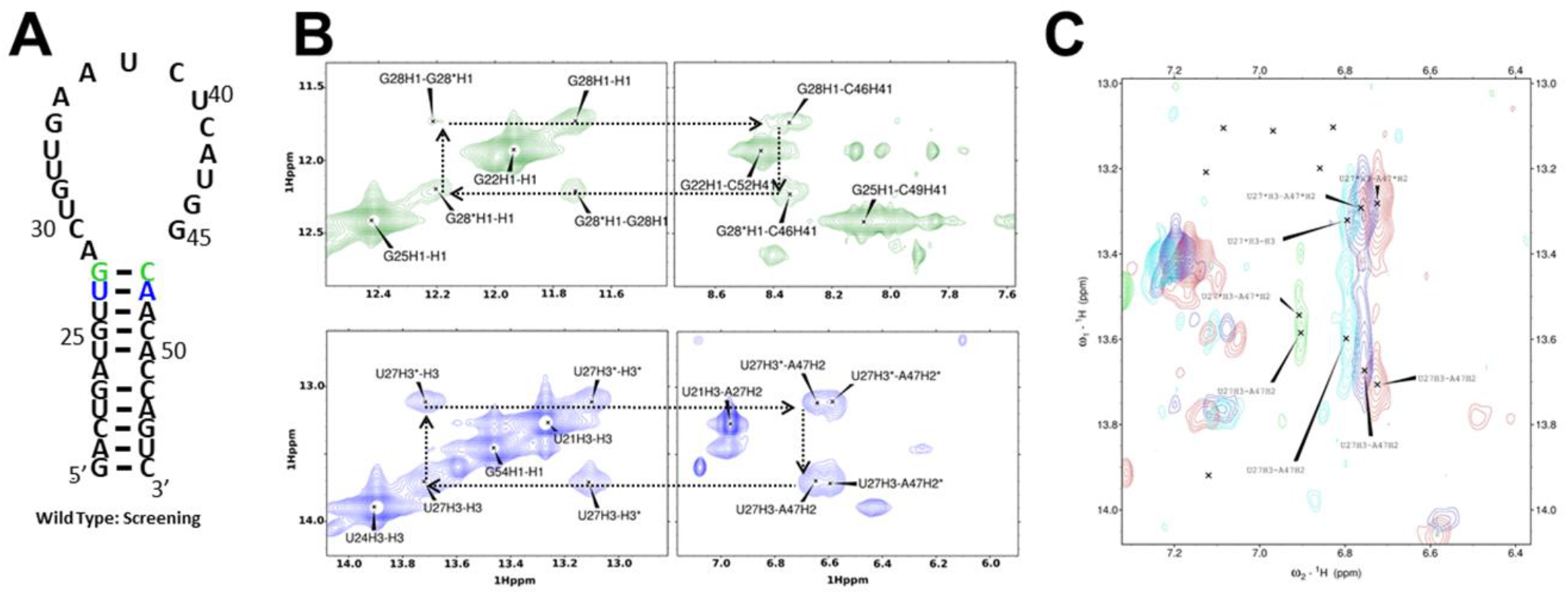
A) Secondary structure of an intermediate (39 nt) pre-miR21 model that fully recapitulates the dynamic behavior at the Dicer cleavage site observed in the actual pre-miR-21 of 59 nts, but has improved NMR properties. B). Sections of ^1^H-^1^H 2D NOESY spectra showing slow chemical exchange for both G28-C46 (green) and U27-A47 (blue). C). Temperature gradient showing progressive coalescence at 37C for the U27-A47 cross peak (Red 5C, Blue 15C, Cyan 25C, Green 37C).

We conclude that the Dicer cleavage site of pre-miR-21 occupies two distinct approximately equally populated conformations in slow exchange (about 10 Hz), corresponding to bulged-out and stacked-in A29, that are not observed in shorter model hairpins used to simplify the NMR analysis. We were intrigued by the possibility that the slow flipping of A29 (Figure 2E) could contribute to inefficient processing of miR-21 by Dicer.

### A29 base flipping modulates miR-21 processing

To explore how changes in sequence and structure affect conformational dynamics of A29, we prepared a series of pre-miR-21 constructs containing specific mutations in the apical loop. These mutations were selected based on previous reports to show unchanged (mutant 1), increased (mutant 2), or decreased (mutant 3) processing efficiencies in cells and in vitro^15^. The secondary structures obtained from the NMR analysis of each mutant are shown in Figure 4A. In all cases, the lower helix is unaffected by the mutations (below the U26-A48 base pair) and the apical loop structures for the mutant sequences remain unpaired, with deviations localized in all cases at or near the mutation site (Figure 4B). In Figure 4C, we show sections of the NOESY spectra collected for the different mutants at 5 °C, highlighting the resonances of U27 and A47. The off-diagonal chemical exchange peaks for U27 observed in the wild-type sequence (Figure 2B) were no longer observed in any of the mutants, but the cross peak between U27 NH and A29 H2 remains visible due to the sharp H2 resonance and provides a useful reporter to monitor the conformation of the Dicer cleavage site.

**Figure 4.**
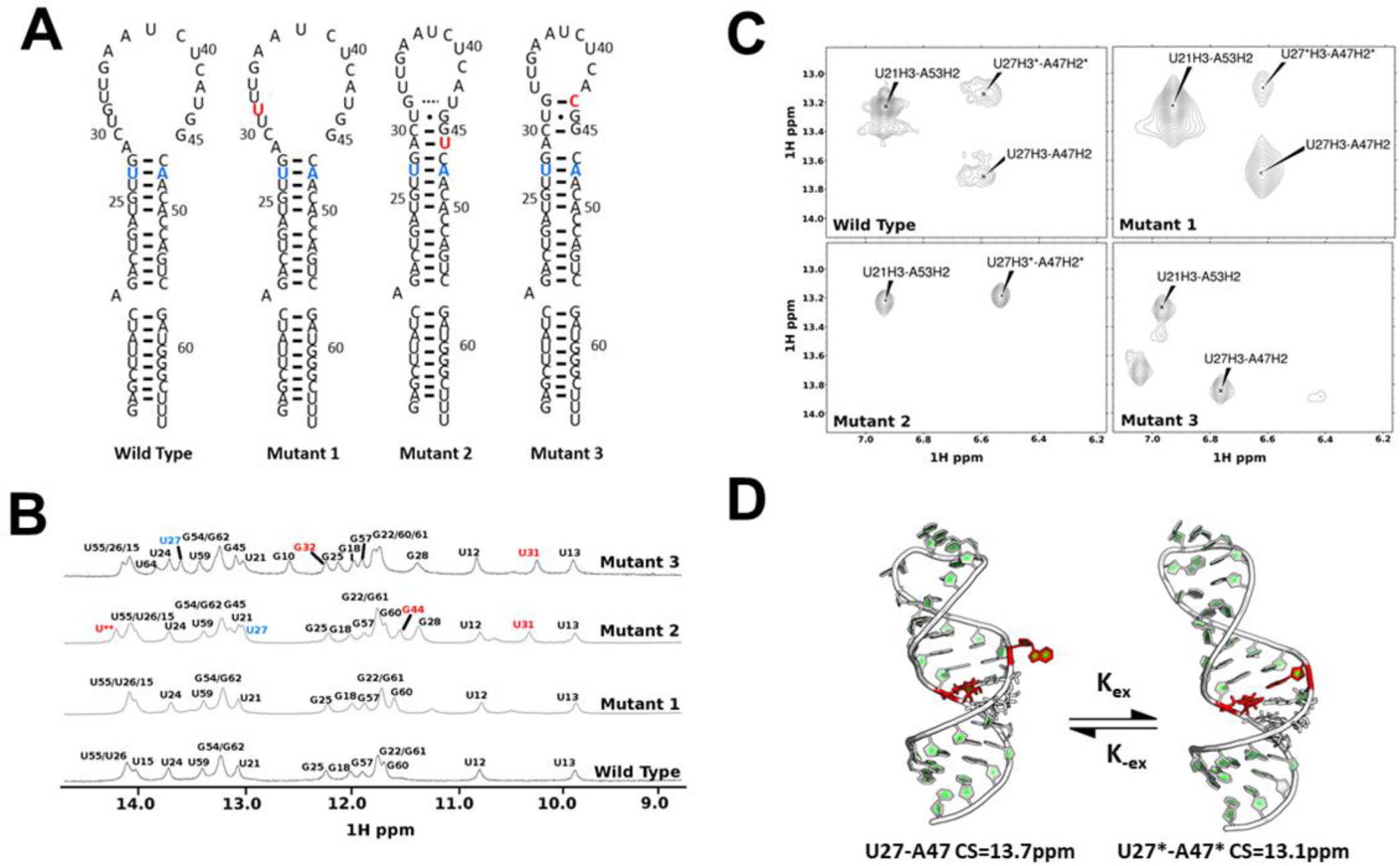
A) Mutant pre-miR-21 sequences used to study the structure and biochemical consequences of conformational exchange at the Dicer cleavage site. B) Systematic analysis of the NMR spectra reveals that the secondary structure of the premiR-21 mutants is unchanged except near the site of the mutation, but (C) mutants 2 and 3 have a restrained upper apical loop conformation and different dynamic behavior. (D) The two conformational states correspond to A29 stacked-in (upfield chemical shift for U27) or A29 bulged-out (downfield U27 chemical shift). Mutant 2 favors the upfield chemical shift (13.1 ppm), corresponding to the A29-in conformation, and has increased processing rate^15^, while mutant 3 exclusively populates the other U27 NH chemical shift (13.8 ppm), corresponding to the A29-out conformation, and has decreased processing instead^15^.

The wild type and mutant 1 pre-miR-21 constructs both lack NH resonances for the proposed tandem UG/GU wobble pairs in the apical loop, or for G28-C46 and C30-G45 (Figure 4B). We also observe two different conformational states on the U27 NH – A47 H2 cross-peak for these two RNAs (Figure 4C). An increase in peak intensity is observed in mutant 1 for the lower chemical shift peak (13.7 ppm)) and a corresponding decrease in peak intensity for the 13.1 ppm conformation, compared to wild-type pre-miR-21. The chemical exchange resonance between these two resonances, which is clearly visible in the wild-type sequence, was not visible in mutant 1 as a result of the altered conformational equilibrium, that now significantly favors the U27-13.7 ppm conformation. The 13.1 ppm chemical shift for U27 is similar to what is observed for the short pre-miR21 construct in Figure 2B, when A29 is stacked inside the RNA helix^31, 34^. This attribution is consistent with the strong ring current associated with stacking of purines inside RNA helices, which could conceivably shift the U27 NH upfield both through the direct ring current of A29, and through stabilization of the G28-C46 base pair.

Mutant 2 places an additional uracil across from A29 leading to formation of a base pair, confirmed by the new NH resonance at 14.2 ppm (Figure 4B). We also observe new NH signals for U31 and G44, suggesting stabilization of the U31-G44 base pair, but the G32-U43 wobble pair remains unstable or open, not observed in the NMR even at 5 °C. The RNA conformation is strongly shifted to the upfield 13.1 ppm conformation (Figure 4C): the peak at 13.7 ppm is no longer observed in mutant 2.

Mutant 3 swaps U43 for C43. This mutation stabilizes the U31:G45 and G32:C43 base pairs, and preferentially selects the lower (13.7 ppm) conformational state (Figure 4C). For mutant 3, we also identify a weak NOE interaction between G28 and G45, although the G45 peak remains broad and weak, as observed in the short mutated pre-miR-21 hairpin (PDB 2MNC), strongly suggestive of a bulged-out conformation for A29^33^. The downfield shift is also consistent with a bulged-out conformation for A29, because the large ring currents which would shift the U27 NH upfield, as observed in mutant 2, would not be present in this conformation. Altogether, these results support the notion that the chemical exchange monitored by the U27-A47 cross peaks corresponds to the flipping in (13.1 ppm chemical shift) and out (13.7 ppm chemical shift) of A29, as modeled in Figure 4D.

Both mutants 2 and 3 restrain the local structure near the Dicer cleavage site compared to the wild-type or mutant 1, by enforcing cross-strand hydrogen bonds in G28 and G45 (Figure 4B). They also alter the local conformational equilibrium, but in opposite ways, as revealed by the spin populations of the U27-A47 cross-peak; mutant 2 favors the peak at 13.1 ppm (A29 stacked-in) while mutant 3 favors the peak at 13.7 ppm (A29 bulged-out). Both mutant 2 and mutant 3 induce the same ‘closed’ conformation of the GU base pairs above A29, yet the two mutants have very different effects on RNA processing efficiency^15^.

We selected these three mutants because they are processed with very different efficiency biochemically and in cells^11, 15, 26^. Mutant 2 is processed more efficiently, while mutant 3 is less efficiently processed. Our data show that mutant 2 adopts the A29-in (U27 13.1 ppm) conformation and is processed relatively efficiently by Dicer-TRBP in vitro^11, 26^. In contrast, mutant 3, which selects the A29-out conformation (U27 13.7 ppm) was shown to be processed inefficiently^15^.

These correlations strongly suggest that it is not closing of the GU base pairs per se, but the conformational state of A29, which determines the processing efficiency. In this model, the ‘off’ or ‘slow’ state would correspond to bulged-out A29 (U27 at 13.7ppm), while the ‘on’ or ‘fast’ conformation corresponds instead to A29 stacked within the helix (U27 at 13.1ppm). This model is consistent with the known mechanism of RNAseIII enzymes^36, 37^, which places a magnesium ion between two phosphate groups prior to in-line cleavage. When A29 is bulged out, proper placement of this magnesium ion would be disfavored. Alternatively, or in addition, bulging out of A29 might disrupt the regular A-form helical structure recognized by the dsRBDs of Dicer and TRBP.

### RNA-binding ligands stabilize the ‘off’ conformational state

While the data presented above support the idea that the conformation of A29 controls processing efficiency, we used several compounds which inhibit processing of pre-miR-21 and bind near the Dicer cleavage site to further test this hypothesis. Namely, we examined the conformational and dynamic consequences of binding of (Figure 5) streptomycin, mitoxantrone and peptide L50, as well as a new macrocyclic peptide designed in house to improve the structural rigidity of the peptide backbone by using paired type 1’ beta turns (KG and NG) compared to the dPro-LPro turn used in L50 (M. Shortridge et al, manuscript in preparation), and with a diamino propionic acid to replace Arg 3 in peptide L50, as was done in a similar project^38^. In each case, we titrated samples of the intermediate (39 nt) pre-miR-21 hairpin with the ligand to investigate whether binding would modulate the dynamic signature of the Dicer cleavage site.

**Figure 5.**
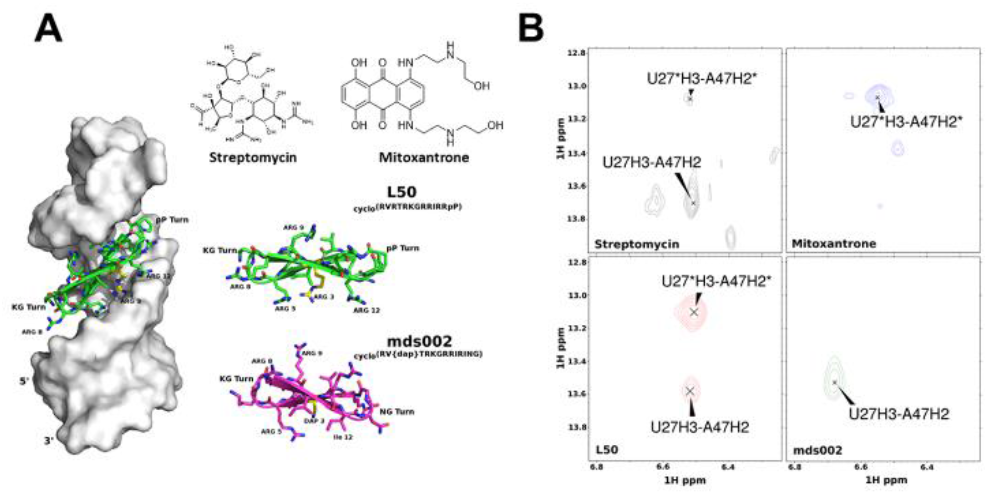
A) Small molecules and cyclic peptides that bind to the pre-miR-21 apical loop modulate its structure. B) Conformational dynamic landscape for the Dicer cleavage site in the presence of different ligands, as monitored by the U27-A47 base pair. Streptomycin and peptide L50 do not significantly alter the conformation of pre-microRNA-21; mitoxantrone induces the stacked-in conformation, but the entire pre-miR-21 spectrum is broadened-out by non-specific binding; peptide mds002 induces the bulged-out conformation and shuts down pre-miR-21 processing (Figure S4).

Streptomycin and L50, which only weakly inhibit pre-miR-21 processing by Dicer in vitro^31^, largely retain the chemical exchange signature of the unbound, wild-type sequence. Mitoxantrone induces the stacked-in A29 conformation represented by the 13.1 ppm chemical shift of U27 (Figure 5B). However, the NOESY spectrum of mitoxantrone-bound premiR21 shows very significant peak broadening, indicative of aggregation and/or multiple binding sites, making it impossible to draw any conclusion; it is clearly a non-specific RNA-binding molecule^39^ whose cellular effect on miR-21 levels^40^ likely originates from cytotoxicity.

In contrast, the new peptide with stabilized turn conformation selects the A29-out state, represented by the 13.7 ppm chemical shift of U27 NH. Consistent with our model, titration of the new peptide into pre-miR-21 completely inhibits processing of pre-miR-21 by Dicer-TRBP, to an extent we have not observed with any other ligand so far (Figure S4). Like peptide L50, this macrocycle binds in the major groove, opposite from Dicer, yet it affects processing even if it cannot interfere sterically with the enzyme. Its effect on the conformation of A29 provides a plausible mechanism for its biochemical activity.

## Discussion

All microRNA precursors share the same stem-loop structures recognized by the processing enzyme complexes (Drosha/DGCR8 and Dicer-TRBP) and have similar binding affinities for them. Yet individual microRNAs are generated with different efficiency and kinetics, leading to different levels of expression in the cell^4, 11, 26, 40^. The structural and/or dynamic features of a pre-miRNA which determine processing efficiency remain unclear. For example, pre-miR-21 is inefficiently processed by Dicer while Let-7 is not^11, 41^; indeed, we also observe very slow processing of pre-miR-21 using in house prepared Dicer-TRBP that is highly active on other pre-miRNAs, e.g. Let-7 (Fig. 1C). This observation suggests miR-21 is normally repressed, to tightly regulate its expression given its role in multiple pathologies and in modulating the immune system^42-44^, but also primed to be up-regulated in response to specific signals (e.g. TGFb, Smad, IL-6/STAT3)^45-47^. Yet, the sequence and secondary structure of pre-miR-21 do not contain any obvious impediment to processing^29^.

We observe a very slow (10 Hz rate) conformational re-arrangement in pre-miR-21 centered on nucleotide A29, at the 5’-Dicer cleavage site which occupies two similarly populated conformations where the base is stacked into the double helix or bulged out. We propose that these two conformations correspond to differently processed structures, similar to a dynamic ‘on’/’off’ or ‘fast’/’slow’ switch, which can be modulated by single nucleotide mutations that affect miR-21 accumulation in vitro and in cells^11, 15, 26^. Mutations that stabilize the marginally stable wobble base pairs near A29 can either increase or decrease Dicer processing, but mutations that stabilize the bulged-out conformation lead to inefficient processing, while mutations that stabilize the stacked-in conformation are processed more efficiently and lead to greater accumulation of miR-21 in cells. To provide further support for the proposed mechanism, we show that the ‘off’ or ‘slow’ conformational state can also be stabilized by cyclic peptides and small molecule ligands that bind in the major groove and would not be expected to sterically or chemically interfere with the processing enzymes which act instead in the minor groove, and yet inhibit processing in vitro^31^.

The conformational exchange is very slow, occurring at a rate of 5-10 Hz, as revealed by the observation of two distinctive signals for U27 and G28 NH in slow exchange on the NMR time scale. In the actual pre-miR-21 of 59 nts, but not in shorter RNA models studied by us and others in the past to simplify the NMR analysis, the U27 and G28 resonances are split; the upfield signal corresponds to A29 stacked-in, and the downfield signal corresponds instead to the bulged-out conformation. These two conformations are only observed when at least a full round of double helix is included in the construct; shorter constructs^34^ adopt instead a single dominant conformation, corresponding to A29 stacked-in and forming a protonated A+-G base pair at least transiently, as revealed by unusually shifted and sharp A29-NH2 resonances (Figure S2)^34^.

Conformational exchange associated with protonation of A29 was reported to occur at a much faster exchange rate (ms), leading to the dynamic formation of the A29+-G45 base pair^34^. However, the model RNA used in that study was much shorter than the complete pre-miRNA that is the Dicer substrate and is not a biologically or biochemically relevant model. In the complete pre-miR-21 that is the Dicer substrate, we no longer observe the characteristic NH2 signal originating from the A29+-G45 base pair (Fig. S2). It is possible that both conformational exchange mechanisms are present to first set up a conformation conducive to binding (slow motion at 0.1s rate), while formation of the protonated base pair could facilitate the catalytic step (faster motion at ms rate^34^).

These results have implications for understanding the mechanism of microRNA processing, and how RNA-binding proteins can regulate processing efficiency by affecting microRNA conformation. Conformational transitions in RNA secondary structure, such as observed here, have slow kinetics and might require the binding energy of RNA-binding proteins to occur in the cell. The behavior we observe for miR-21 is not unique; we observe the same conformational exchange in another cancer-causing microRNA (Figure S5), miR-10b, which is generally repressed and only expressed in metastatic cancer cell lines^7, 48^. Since small molecule and peptidic ligands can affect the RNA conformation, our results suggest a strategy to inhibit micro-RNA maturation by pushing the RNA into the ‘off’ or ‘slow’ conformation, or activate the expression of tumor suppressor microRNAs by favoring population of the ‘on’ or ‘fast’ state instead (Figure S6).

## Materials and Methods

### RNA sample preparation

All RNAs were prepared in house by *in vitro* transcription, using well-established methods^49^. Briefly, RNA transcription and purification used DNA oligonucleotide templates (IDT) and T7 RNA polymerase purified *in house* using affinity capture of a His-tagged protein. 1 μL of 8 μM TOP DNA (5’-CTATAGTGAGTCGTATTA-3’), corresponding to the phage T7 RNA polymerase promoter region, was annealed to 80 μL of 100 μM template sequences with 13 mM MgCl2, heated to 95 °C for 4 min, then allowed to anneal to room temperature over 20 min. The annealing mixture was incubated with 5 mM of each of the four NTPs (ATP, GTP, UTP and CTP, from Sigma), 1x transcription buffer, 8% PEG-8000 and 35 mM magnesium chloride with 0.4 mg/mL T7 RNA polymerase.

All RNA samples were purified from crude transcriptions by 20% denaturing polyacrylamide gel electrophoresis (PAGE), electroeluted and concentrated by ethanol precipitation. The samples were re-dissolved in 12 mL of high salt wash (700 mM NaCl, 200 mM KCl, in 10 mM potassium phosphate at pH 6.5, with 10 µM EDTA), concentrated using Centriprep conical concentrators (3,000 Da MWC, Millipore). The RNA was then slowly exchanged into low salt storage buffer (10 mM potassium phosphate at pH 6.5, with 10 mM NaCl and 10 µM EDTA). Prior to NMR experiments, all RNA samples were finally desalted using NAP-10 gravity columns, lyophilized and re-dissolved in buffer, then annealed by heating for 4 min to 90 °C and snap cooling at -20 °C. For experiments investigating non-exchangeable protons, samples were lyophilized to dryness and dissolved into 99.99% D2O. Samples used to study exchangeable protons were dissolved instead in 95% H2O/5% D2O.

### Small molecules and peptide synthesis

Small molecules were purchased in the highest purity available from Sigma and used without further purification. Peptides were prepared by Fmoc chemistry, as previously described^31, 38, 50^. Coupling reactions were completed per manufacturer protocol, purified by HPLC; purity and structure were confirmed by NMR.

### NMR spectroscopy and Data Analysis

NMR data were collected on Bruker 500, 600 and 800 MHz spectrometers equipped with HCN cryogenic probes. Experiments were conducted at multiple temperatures using the 1D ^1^H excitation sculpting pulse sequence for water suppression. Chemical exchange results were collected from 2D ^1^H-^1^H nuclear Overhauser effect spectroscopy (NOESY) spectra, collected in H2O NMR buffer (20 mM potassium phosphate pH 6.5, 50 mM sodium chloride) at 5 °C. NOESY spectra with mixing times of 10 ms and 100 ms were recorded to measure exchange rates with EXSYCALC (Mestrenova). All NMR data were processed using Bruker Topspin (3.1) or NMRPipe and visualized with Topspin and Sparky.

### Dicer/TRBP

The Dicer/TRBP enzyme complex was expressed and purified *in house* using published methods^11, 20^ and stored at -80 °C in 20 μL aliquots until needed. The gene encoding human full-length Dicer was inserted into a modified pFastBac1 vector (Invitrogen), which has an N-terminal glutathione-S-transferase (GST) tag. The GST fusion proteins were expressed in HiFive insect cells following standard procedures and isolated by glutathione affinity chromatography using buffer A (20 mM Tris-HCl pH 8.0, 150 mM NaCl, 2 mM DTT). After on-column cleavage by Tobacco Etch Virus (TEV) protease overnight, the proteins were eluted, concentrated and loaded onto a Superdex 200 10/300 GL column per final size-exclusion purification. Sequences coding for human full-length TARBP2 were cloned into a modified pET28a vector with an N-terminal GB1 tag. The clones were transformed into Rosetta 2 cells and the transformants were grown in LB media at 37 °C until the cells reached a density corresponding to an OD reading of 0.8 at 600 nm. Protein expression was induced by addition of 0.2 mM IPTG at 18 °C for 20 hours. The cells were harvested, pelleted and resuspended in 20 mM Tris-HCl pH 8.0, 500 mM NaCl, 25 mM imidazole, with 5 mM β-ME. Cells were lysed by sonication on ice then pelleted by centrifugation. The crude lysate was applied to a nickel affinity column and eluted in 20 mM Tris-HCl pH 8.0, 500 mM NaCl, 250 mM imidazole buffer, with 5 mM β-ME. TEV protease was added to remove the His-GB1 tag at 4 °C overnight. The sample was concentrated and loaded onto a Superdex 200 10/300 GL column. The purified Dicer and TARBP2 proteins were mixed at a molar ratio of 1:1.3 and loaded onto Superdex 200 10/300 GL column for final purification. The fractions containing the complex were pooled and concentrated to ∼250 nM, then flash frozen for further use in processing assays.

### Pre-miRNAs

All pre-miRNAs were chemically synthesized by Integrated DNA Technologies (IDT) for efficient 5’-end labeling, which was done using a modified T4 PNK reaction. Radiolabeling reactions were carried out in 30 μL volumes, containing 11 μL DNAse/RNAse free water, 3 μL 10x T4 PNK buffer, 3 μL of 10 μM pre-miRNA (IDT), 3 μL of 10 units/uL T4 PNK, 10 μL γ-32P ATP (6,000 Ci/mmol, 150 mCi/mL). Reactions proceeded for 1 hr at 37 °C, followed by the addition of 50 mM EDTA to quench the reaction, along with heating to 95 °C for 10 min to inactivate the enzyme. After the PNK labeling reaction was completed, excess unincorporated nucleotides were removed using Zymo Research oligo clean up columns and RNA was eluted in 15 μL of RNAse/DNAse free water. An equal volume of 2x formamide gel loading buffer was added to the eluted RNA, followed by incubation at 95 °C for 5 min, then loaded onto 20% 19:1 denaturing (8 M urea) polyacrylamide gels. Gels were run for 1.25 hrs at 10 W, covered with plastic wrap and exposed to x-ray film for 30 sec to visualize the labeled RNA. The developed film was laid on the gel and oriented by internal phosphorescent paint positional controls placed near dye molecular weight markers. The area around the developed bands corresponding to the molecular weight of the target RNAs was excised and RNA was extracted from the gel slices by using Zymo Research ZR-small-RNA PAGE recovery kit, eluted into 15 μL of water and stored at -20 °C. These fresh 15 μL samples had a specific reactivity of 200 k CPM, as measured by a Geiger counter with an approximate RNA concentration of 2 μM.

### Enzymatic assays

Dicer/TRBP assays were run immediately (within 2 weeks) after preparation of the labeled RNA to maintain high levels of radioactivity, which are needed because miR-21 processing is inefficient^11, 26^ and high sensitivity is required for accurate quantitation of initial processing rates. Assays were conducted in 60 μL total volume with 25 nM RNA, 1 nM Dicer/TRBP in 1x reaction buffer (20 mM Tris at pH 7.5, 25 mM NaCl, 5 mM MgCl2, 1 mM DTT and 1% glycerol), and in 96-well format in a PCR thermocycler in 0.2 mL PCR well plates to maintain constant temperature and improve reproducibility. This approach allows for direct comparison for different pre-miRNAs along with technical duplication to ensure data reproducibility with regard to enzyme aliquots. Each reaction well (column 1 and/or 2) contained 25 μL of RNA substrate (typical stock RNA concentration was 67 nM but this concentration was titrated for kinetics work). Each enzyme well (column 3) was filled with 50 μL of freshly diluted 2 nM Dicer-TRBP while each ligand well (column 4) was filled with 50 μL of ligand at different concentrations (0-150 μM). Finally, 20 μL of reaction stop buffer (2x RNA load buffer, 95% formamide, 18 mM EDTA, 0.025% SDS, 0.1% xylene cyanol and 0.1% bromophenol blue) was added to the remaining wells (columns 5-12).

Prior to initiating the reactions, 10 μL of ligand, when relevant, were added into the RNA wells using a multichannel pipette and allowed to equilibrate at 4 °C for 30 min. The temperature of the thermocycler was then raised to 37 °C and samples were incubated for 15 min to reach thermal equilibrium. Reactions were initiated by adding 25 μL of Dicer-TRBP to the RNA substrate wells (with or without ligand). The final enzyme concentration for all experiments was fixed to 1 nM while the final RNA and ligand concentrations were set at 10 µM. The reactions were stopped by removing 5 μL from the reaction well and adding it to stop buffer wells at the indicated time points. Reactions were resolved by 20% 19:1 denaturing (8 M urea) polyacrylamide gel and visualized using a Typhoon Gel imaging system (GE). Bands were quantified in ImageJ and results visualized as ratios of cleaved and uncleaved RNAs, after subtracting the background.

## Supporting Information

Figures S1-S6: 1D ^1^H NMR spectra comparing imino regions for short and full-length pre-miRNA-21 transcripts (Figure S1); NOESY spectra of pre-miR21 stem-loop and corresponding ‘short’ model construct (Figure S2); NOESY spectra probing for conformational switching in the three miR-21 constructs studied (Figure S3); Gel shift assays showing the effect of peptide mds002 on Dicer-TRBP assays for pre-miR-21, and NOESY spectra illustrating effects of peptide mds002 on the pre-miR-21 conformation (Figure S4); NOESY spectra illustrating conformational states and exchange in pre-miR-10b and pre-miR-21 (Figures S5, S6).

## Author Contributions

MDS collected and analyzed the data, synthesized peptides and conducted enzymatic assays, as did MW. WY prepared Dicer-TRBP. The research was conceived by MDS and GV and supervised by GV. The manuscript was written primarily by MDS, MW and GV. All authors have approved the final version of the manuscript.

## Acknowledgements

We wish to thank all members of the Varani group for discussion and support.

## References

1. Kim, V. N., MicroRNA biogenesis: Coordinated cropping and dicing. Nature Reviews Molecular Cell Biology 2005, 6 (5), 376–385.

2. Winter, J.; Jung, S.; Keller, S.; Gregory, R. I.; Diederichs, S., Many roads to maturity: microRNA biogenesis pathways and their regulation. Nat Cell Biol 2009, 11 (3), 228–234.

3. Krol, J.; Loedige, I.; Filipowicz, W., The widespread regulation of microRNA biogenesis, function and decay. Nat Rev Genet 2010, 11 (9), 597–610.

4. Reichholf, B.; Herzog, V. A.; Fasching, N.; Manzenreither, R. A.; Sowemimo, I.; Ameres, S. L., Time-Resolved Small RNA Sequencing Unravels the Molecular Principles of MicroRNA Homeostasis. Molecular Cell 2019, 75 (4), 756–768.

5. Calin, G. A.; Sevignani, C.; Dumitru, C. D.; Hyslop, T.; Noch, E.; Yendamuri, S., Human microRNA genes are frequently located at fragile sites and genomic regions involved in cancers. Proc Natl Acad Sci USA 2004, 101, 2999-3004.

6. Adams, B. D.; Parsons, C.; Walker, L.; Zhang, W. C.; Slack, F. J., Targeting noncoding RNAs in disease. Journal of Clinical Investigation 2017, 127 (3), 761–771.

7. Sheedy, P.; Medarova, Z., The fundamental role of miR-10b in metastatic cancer. American Journal of Cancer Research 2018, 8 (9), 1674–1688.

8. Winkle, M.; El-Daly, S. M.; Fabbri, M.; Calin, G. A., Noncoding RNA therapeutics - challenges and potential solutions. Nature Reviews Drug Discovery 2021, 20 (8), 629–651.

9. Zhang, S.; Chen, L.; Jung, E. J.; Calin, G. A., Targeting MicroRNAs With Small Molecules: From Dream to Reality. Clin Pharmacol Ther 2010, 87 (6), 754–758.

10. Warner, K. D.; Hajdin, C. E.; Weeks, K. M., Principles for targeting RNA with drug-like small molecules. Nature Reviews Drug Discovery 2018, 17 (8), 547–558.

11. Lee, H. Y.; Doudna, J. A., TRBP alters human precursor microRNA processing in vitro. Rna 2012, 18 (11), 2012–2019.

12. Zeng, Y.; Cullen, B. R., Efficient processing of primary microRNA hairpins by drosha requires flanking nonstructured RNA sequences. Journal of Biological Chemistry 2005, 280 (30), 27595–27603.

13. Lee, Y.-F.; Ahn, C.; Han, J.; Choi, H.; Kim, J.; Yim, J.; Lee, J.; Provost, P.; Radmark, O.; Ki, S.; Kim, V. N., The Nuclear RNAseIII Drosha Initiates MicroRNA Processing. Nature 2003, 425, 415–419.

14. Gregory, R. I.; Yan, K.; Amuthan, G.; Cendrimada, T.; Doratotaj, B.; Cooch, N.; Shiekhattar, R., The Microprocessor complex mediates the genesis of microRNAs. Nature 2004, 432, 235–240.

15. Zeng, Y.; Yi, R.; Cullen, B. R., Recognition and cleavage of primary microRNA precursors by the nuclear processing enzyme Drosha. Embo Journal 2005, 24 (1), 138–148.

16. Han, J. J.; Lee, Y.; Yeom, K. H.; Nam, J. W.; Heo, I.; Rhee, J. K.; Sohn, S. Y.; Cho, Y. J.; Zhang, B. T.; Kim, V. N., Molecular basis for the recognition of primary microRNAs by the Drosha-DGCR8 complex. Cell 2006, 125 (5), 887–901.

17. Nguyen, T. A.; Jo, M. H.; Choi, Y. G.; Park, J.; Kwon, S. C.; Hohng, S.; Kim, V. N.; Woo, J. S., Functional Anatomy of the Human Microprocessor. Cell 2015, 161 (6), 1374–1387.

18. Kwon, S. C.; Nguyen, T. A.; Choi, Y. G.; Jo, M. H.; Hohng, S.; Kim, V. N.; Woo, J. S., Structure of Human DROSHA. Cell 2016, 164 (1-2), 81–90.

19. Kwon, S. C.; Baek, S. C.; Choi, Y. G.; Yang, J.; Lee, Y. S.; Woo, J. S.; Kim, V. N., Molecular Basis for the Single-Nucleotide Precision of Primary microRNA Processing. Molecular Cell 2019, 73 (3), 505–518.

20. Liu, Z. M.; Wang, J.; Cheng, H.; Ke, X.; Sun, L.; Zhang, Q. C.; Wang, H. W., Cryo-EM Structure of Human Dicer and Its Complexes with a Pre-miRNA Substrate (vol 173, pg 1191, 2018). Cell 2018, 173 (6), 1549–1550.

21. Zhang, X. X.; Zeng, Y., The terminal loop region controls microRNA processing by Drosha and Dicer. Nucleic Acids Research 2010, 38 (21), 7689–7697.

22. Castilla-Liorente, V.; Nicastro, G.; Ramos, A., Terminal loop-mediated regulation of miRNA biogenesis: selectivity and mechanisms. Biochemical Society Transactions 2013, 41, 861–865.

23. Gu, S.; Jin, L.; Zhang, Y.; Huang, Y.; Zhang, F. J.; Valdmanis, P. N.; Kay, M. A., The Loop Position of shRNAs and Pre-miRNAs Is Critical for the Accuracy of Dicer Processing In Vivo. Cell 2012, 151 (4), 900–911.

24. Michlewski, G.; Guil, S.; Semple, C. A.; Caceres, J. F., Posttranscriptional Regulation of miRNAs Harboring Conserved Terminal Loops. Molecular Cell 2008, 32 (3), 383–393.

25. Michlewski, G.; Caceres, J. F., Antagonistic role of hnRNP A1 and KSRP in the regulation of let-7a biogenesis. Nature Structural & Molecular Biology 2010, 17 (8), 1011–1018.

26. Chakravarthy, S.; Sternberg, S. H.; Kellenberger, C. A.; Doudna, J. A., Substrate-Specific Kinetics of Dicer-Catalyzed RNA Processing. Journal of Molecular Biology 2010, 404 (3), 392–402.

27. Nicastro, G.; Garcia-Mayoral, M. F.; Hollingworth, D.; Kelly, G.; Martin, S. R.; Briata, P.; Gherzi, R.; Ramos, A., Noncanonical G recognition mediates KSRP regulation of let-7 biogenesis. Nature Structural & Molecular Biology 2012, 19 (12), 1282–1286.

28. Trabucchi, M.; Briata, P.; Garcia-Mayoral, M.; Haase, A. D.; Filipowicz, W.; Ramos, A.; Gherzi, R.; Rosenfeld, M. G., The RNA-binding protein KSRP promotes the biogenesis of a subset of microRNAs. Nature 2009, 459 (7249), 1010–1014.

29. Sperber, H.; Beem, A.; Shannon, S.; Jones, R.; Banik, P.; Chen, Y.; Ku, S.; Varani, G.; Yao, S.; Ruohola-Baker, H., miRNA sensitivity to Drosha levels correlates with pre-miRNA secondary structure. RNA 2014, 20, 621–631.

30. MacRae, I. J.; Zhou, K.; Li, F.; Repic, A.; Brooks, A. N.; Cande, W. Z.; Adams, P. D.; Doudna, J. A., Structural Basis for Double-Stranded RNA Processing by Dicer. Science 2006, 311, 195–198.

31. Shortridge, M. D.; Walker, M. J.; Pavelitz, T.; Chen, Y.; Yang, W.; Varani, G., A Macrocyclic Peptide Ligand Binds the Oncogenic MicroRNA-21 Precursor and Suppresses Dicer Processing. ACS Chemical Biology 2017, 12 (6), 1611–1620.

32. Chen, Y.; Yang, F.; Zubovic, L.; Pavelitz, T.; Yang, W.; Godin, K.; Walker, M.; Zheng, S.; Macchi, P.; Varani, G., Targeted inhibition of oncogenic miR-21 maturation with designed RNA-binding proteins. Nat Chem Biol 2016, 12, 717–723.

33. Chirayil, S.; Wu, Q.; Amezcua, C.; Luebke, K. J., NMR Chracterization of an Oligonucleotide Model of the MiR21 Pre-Element. PLOS one 2014, 9 (9), e108231.

34. Baisden, J. T.; Boyer, J. A.; Zhao, B.; Hammond, S. M.; Zhang, Q., Visualizing a protonated RNA state that modulates microRNA-21 maturation. Nature Chemical Biology 2021, 17 (1), 80–88.

35. Goddard, T. D.; Kneller, D. G., Sparky 3. University of California, San Francisco.

36. Blaszczyk, J.; Tropea, J. E.; Bubunenko, M.; Routzahn, K. M.; Waugh, D. S.; Court, D. L.; Ji, X., Crystallographic and modeling studies of RNase III suggest a mechanism for double-stranded RNA cleavage. Structure 2001, 9, 1225–1236.

37. Zhang, H.; Kolb, F. A.; Jaskiewicz, W., Single Processing Center Models for Human Dicer and Bacterial RNase III. Cell 2004, 118, 57–68.

38. Shortridge, M. D.; Wille, P. T.; Jones, A. N.; Davidson, A.; Bogdanovic, J.; Arts, E.; Karn, J.; Robinson, J. A.; Varani, G., An ultra-high affinity ligand of HIV-1 TAR reveals the RNA structure recognized by P-TEFb. Nucleic Acids Res 2019, 47 (3), 1523–1531.

39. Liu, Y.; Peacey, E.; Dickson, J.; Donahue, C. P.; Zheng, S.; Varani, G.; Wolfe, M. S., Mitoxantrone analogues as ligands for a stem-loop structure of tau pre-mRNA. Journal of medicinal chemistry 2009, 52 (21), 6523–6526.

40. Velagapudi, S. P.; Costales, M. G.; Vummidi, B. R.; Nakai, Y.; Angelbello, A. J.; Tran, T.; Haniff, H. S.; Matsumoto, Y.; Wang, Z. F.; Chatterjee, A. K.; Childs-Disney, J. L.; Disney, M. D., Approved Anti-cancer Drugs Target Oncogenic Non-coding RNAs. Cell Chemical Biology 2018, 25 (9), 1086-1094.e7.

41. Feng, Y.; Zhang, X. X.; Graves, P.; Zeng, Y., A comprehensive analysis of precursor microRNA cleavage by human Dicer. Rna-a Publication of the Rna Society 2012, 18 (11), 2083–2092.

42. Sheedy, F. J., Turning 21: induction of miR-21 as a key swith in the inflammatory response. Frontiers in Immunology 2015, 6 (19) doi: 10.3389/fimmu.2015.00019. eCollection 2015.

43. Lu, T. X.; Hartner, J.; Lim, E. J.; Fabry, V.; Mingler, M. K.; Cole, E. T.; Orkin, S. H.; Aronow, B. J.; Rothenberg, M. E., MicroRNA-21 Limits In Vivo Immune Response-Mediated Activation of the IL-12/IFN-gamma Pathway, Th1 Polarization, and the Severity of Delayed-Type Hypersensitivity. Journal of Immunology 2011, 187 (6), 3362–3373.

44. Medina, P. P.; Nolde, M.; Slack, F. J., OncomiR addiction in an in vivo model of microRNA-21-induced pre-B-cell lymphoma. Nature 2010, 467, 86–90.

45. Li, Q.; Zhang, D. X.; Wang, Y. B.; Sun, P.; Hou, X. D.; Larner, J.; Xiong, W. J.; Mi, J., MiR-21/Smad 7 signaling determines TGF-beta 1-induced CAF formation. Scientific Reports 2013, 3, Article number: 2038.

46. Iliopoulos, D.; Jaeger, S. A.; Hirsch, H. A.; Bulyk, M. L.; Struhl, K., STAT3 Activation of miR-21 and miR-181b-1 via PTEN and CYLD Are Part of the Epigenetic Switch Linking Inflammation to Cancer. Molecular Cell 2010, 39 (4), 493–506.

47. Davis, B. N.; Hilyard, A. C.; Lagna, G.; Hata, A., SMAD proteins control DROSHA-mediated microRNA maturation. Nature 2008, 454, 56–61.

48. Ma, L.; Teruya-Feldstein, J.; Weinberg, R. A., Tumour invasion and metastasis initiated by microRNA 10b in breast cancer. Nature 2007, 449, 682–688.

49. Varani, G.; Aboul-ela, F.; Allain, F. H.-T., NMR Investigations of RNA Structure. Progr. NMR Spectr. 1996, 29, 51–127.

50. Davidson, A.; Patora-Komisarska, K.; Robinson, J. A.; Varani, G., Essential structural requirements for specific recognition of HIV TAR RNA by peptide mimetics of Tat protein. Nucleic Acids Research 2011, 39, 248–256.

