## Supplementary material for "A slow dynamic RNA switch regulates processing of microRNA-21"

+ 1 206 543 7113

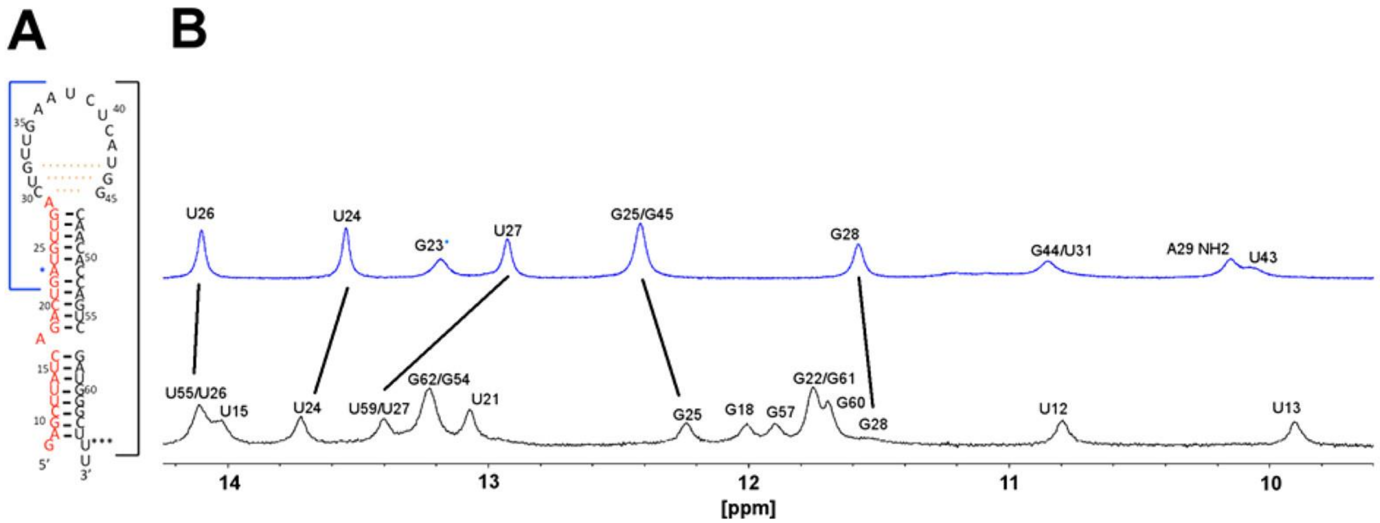


**Figure S1**. **Pre-miRNA-21 stem-loops are characterized by 'open' apical loop structures.** (**A**) Secondary structure of the pre-miR21 stem-loop with the mature miR-21 5p sequence in red. The full-length pre-miR-21 is shown and boxed with black outline; G8 and U65*** were swapped, compared to the wild-type sequence, to improve *in vitro* transcription yields for NMR analysis. The small hairpin model is boxed with a blue line; A23 was switched to G23* to improve *in vitro* transcription yields. Dashes identify weak base pairs observed only at low temperature (5C), but not at 37C. (**B**) The 1D $^1$H imino region for both the short transcript (top, blue) and the full length (bottom, black) pre-miR-21 show close correspondence of equivalent signals (black line), indicating identical folding; both transcripts lack NH signals from the apical loop, which is unpaired and highly flexible at physiological temperatures.

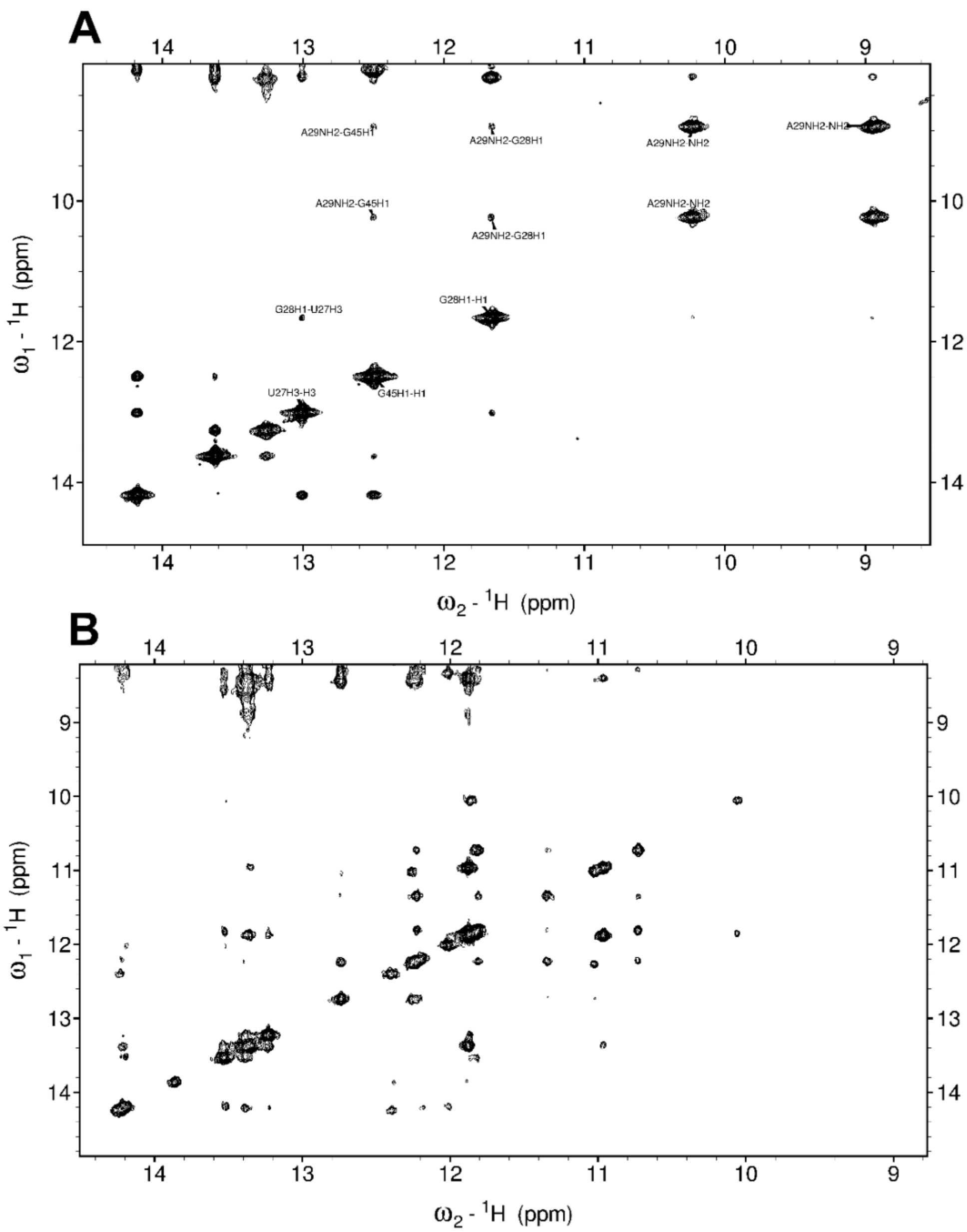


**Figure S2**. **NOESY spectra recorded at 5ºC and 100ms mixing time for both the 'short' stem-loop model and the actual pre-miR21 stem-loop.** (**A**) The very distinctive amino signals for the protonated A29 base NH2s are clearly visible at approximately 8.9 and 10.1ppm, with clear NOE interactions to both G28 and G45 imino protons that facilitate their unambiguous assignment. (**B**) For the full pre-miR-21 sequence, these amino signals are no longer visible (the resonance near 10.0 ppm is from the U13:G60 pair in the lower part of the helix). Neither of the tandem putative apical loop GU wobble pairs (U31:G44 or G32:U43) are visible in the short or full-length sequence.

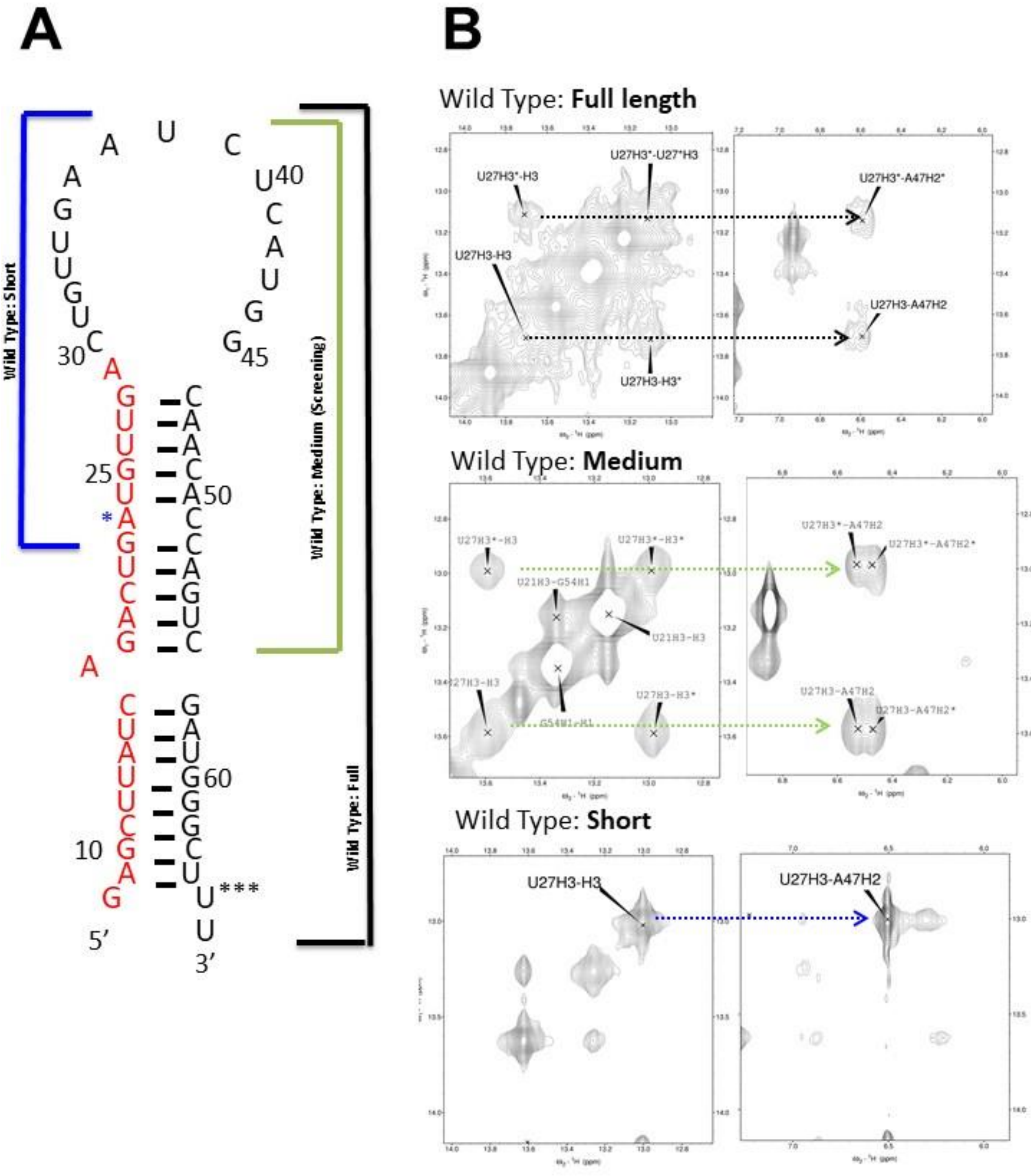
A
B
Wild Type: Short
Wild Type: Medium (Screening)
Wild Type: Full
5′
3′
Wild Type: Full length
U27H3*-H3
U27H3*-U27*H3
U27H3-H3
U27H3-H3*
U27H3*-A47H2*
U27H3-A47H2
Wild Type: Medium
U27H3*-H3
U27H3*-H3*
U21H3-G54H1
U21H3-H3
U27H3-H3
U27H3-H3*
G54H1-H1
U27H3*-A47H2
U27H3*-A47H2*
U27H3-A47H2
U27H3-A47H2*
Wild Type: Short
U27H3-H3
U27H3-A47H2

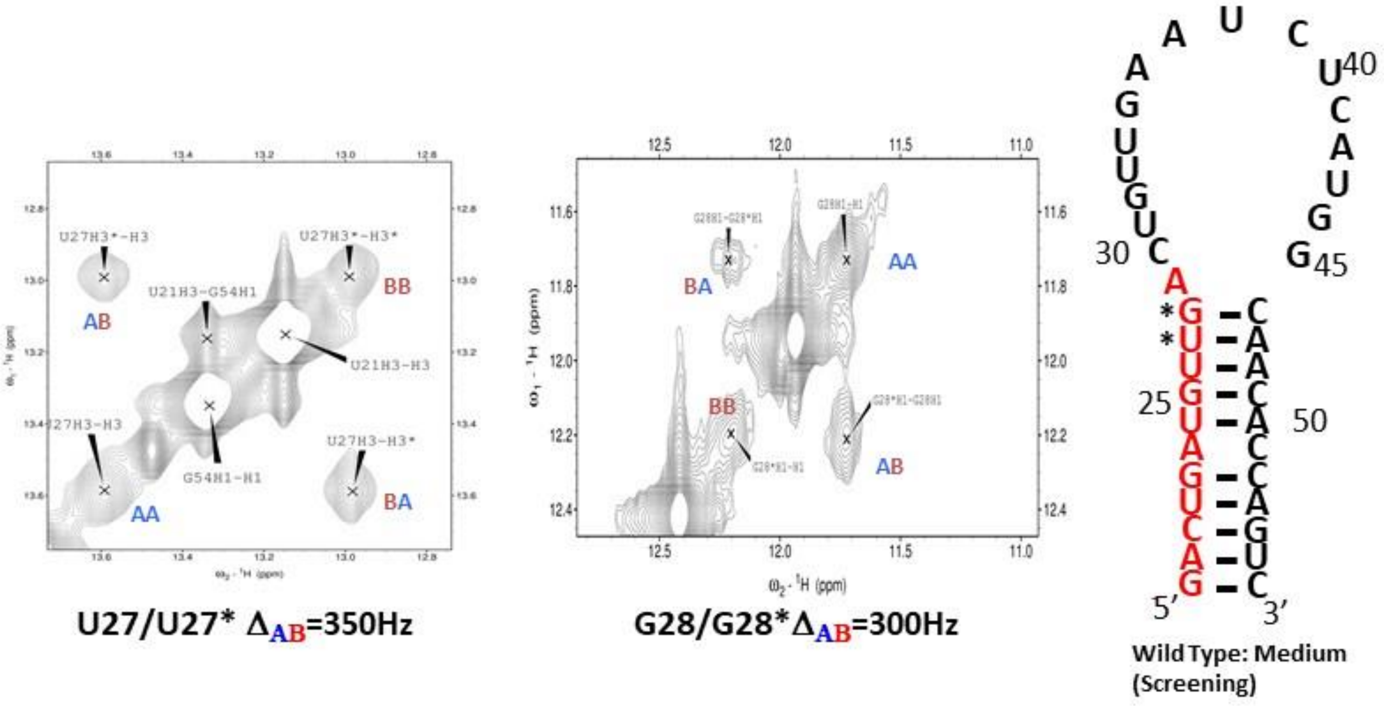


**Figure S3. A slow conformational switch at the Dicer cleavage site of pre-miR-21.** (**A**) The three miR-21 sequences used in this study. Short miR-21 (nts G22-C52); intermediate (nts G18-56); and full pre-miR-21 (nts G8-U66); this last RNA corresponds to the complete pre-miR-21 sequence. For the short RNA sequence, an A23G switch was used to improve transcription (*); similarly, the G8-U65 pair (***) was swapped to CG to improve transcription in the full pre-miR-21. (**B**) All spectra were collected at 800 MHz, at 0.5mM RNA concentration in 10mM phosphate, 10mM NaCl, 100μM EDTA pH 6.5 and at 5 C. A conformational switch is observed in the full length pre-miR-21 transcript in which two U27 NH resonances have an NOE interaction with A47 H2, corresponding to two distinct structural states of the pre-miRNA. The short transcript shown at the bottom does not reproduce the dynamic switch observed in the complete pre-miR-21 RNA hairpin, while the intermediate construct does so. In this intermediate construct, the improved spectral quality allows observation of two chemical shifts for A27H2 as well, which are also visible, but less clearly, in the spectra of the full length pre-miR-21. (**C**) Chemical exchange of U27 and G28 in the intermediate pre-miR-21 construct (shown on the right) that reproduces the behavior of full pre-miR-21. The reduced size of this intermediate pre-miR-21 model leads to improved line width and spectral resolution. Unlike the short sequence, this model accurately reproduces the dynamic behavior found in the complete pre-miR-21, as shown in the 2D NMR spectra for the exchange peaks of U27 and allows observation of exchange peaks for G28 as well, close to the diagonal.

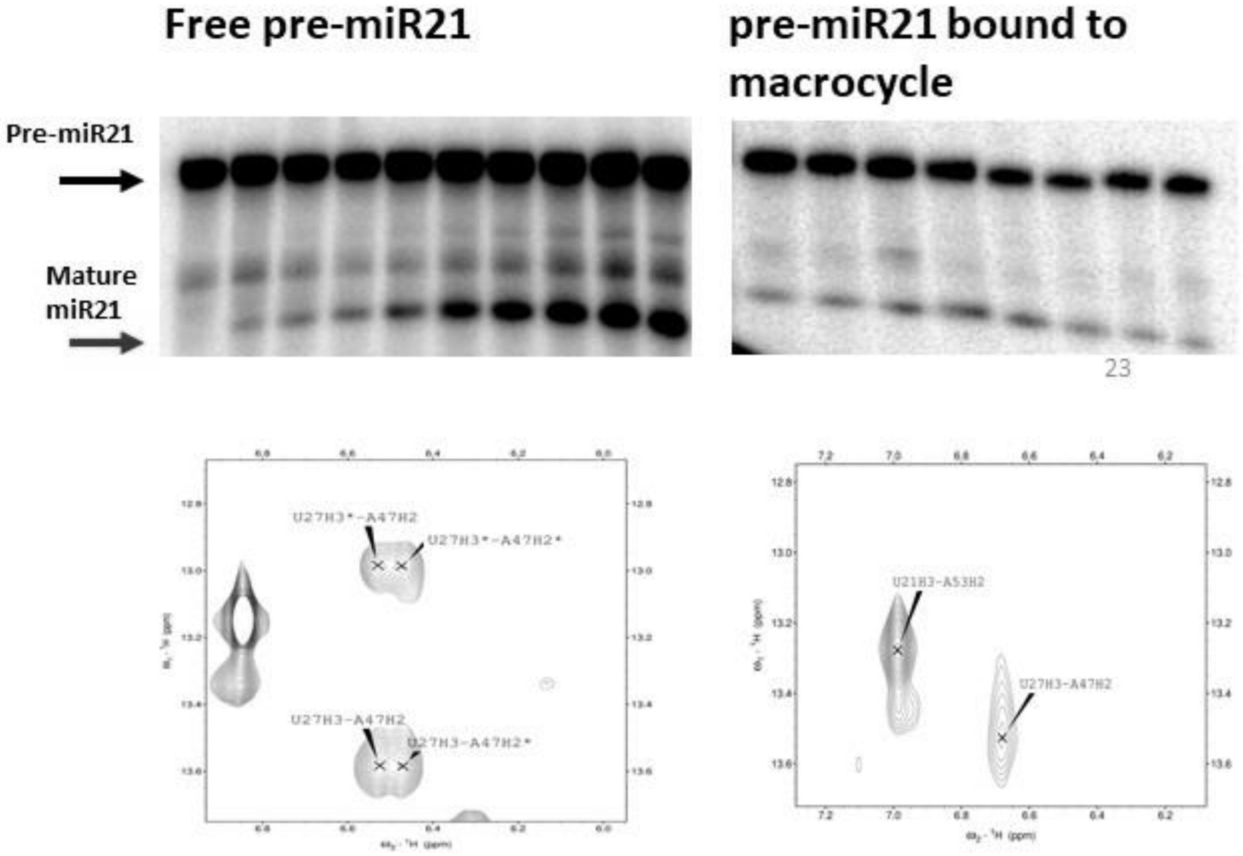


**Figure S4. Inhibition of Dicer-TRBP correlates with selection of the A29-out conformation.** Processing of wild type pre-miR-21 is inherently slow and inefficient (left). However, compound mds002, which binds to pre-miR-21 with high affinity, strongly further inhibits processing of pre-miR-21 in Dicer-TRBP assays when 10 μM peptide is present (right). The peptide selects for the A29-out conformation (bottom, right), as indicated by the U27 NH chemical shift value of 13.6ppm, while A29 occupies two conformational states (A29-in and A29-out, bottom, left) in the absence of the peptide.

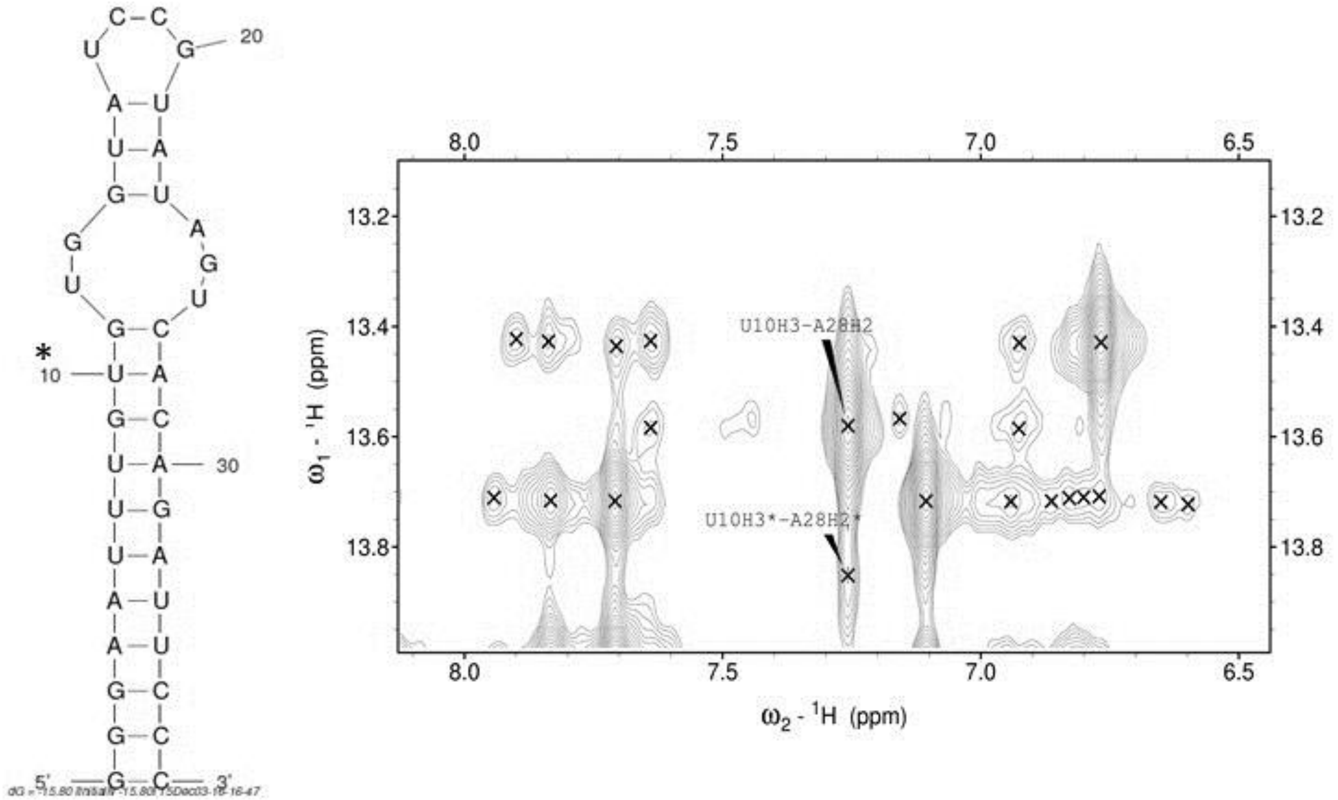


**Figure S5. Slow conformational dynamics in pre-miR-10b.** Slow dynamic exchange signals for pre-miR-10b, whose secondary structure is shown on the left. A section of the NOESY spectrum of pre-miR-10b recorded at 5C identifies chemical exchange peaks corresponding to the U10-A28 base pair. Similar to what is observed for pre-miR-21, two peaks are observed for U10 NH, corresponding to the UA base pair indicated by an asterisk, as a result of conformational exchange in the asymmetric internal loop above it.

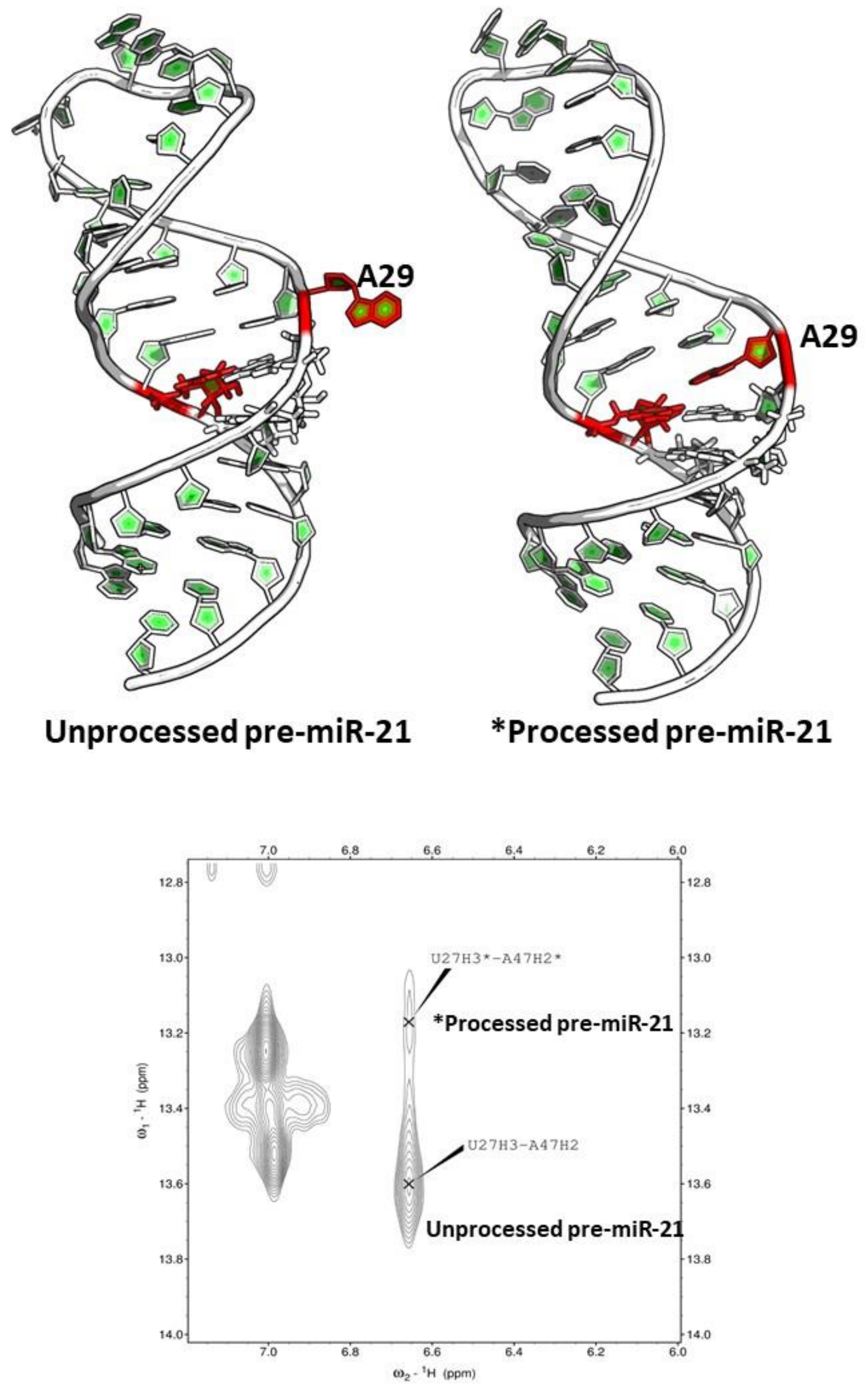


**Figure S6. The NMR investigation of pre-miR-21 identified two conformational states for A29, which correspond to A29 bulged out or stacked into the helix.** The dynamic switch involving A29, located at the 5'-Dicer cleavage site, modulates pre-miR-21 processing. In the slowly processed 'off' state, A29 is bulged out (left); when A29 is stacked in (right), pre-miR-21 is more efficiently processed in its 'on' state.
